# Intersectional roles of trait covariation and phenotypic plasticity in the coloration and behavior of an African cichlid

**DOI:** 10.1101/2025.03.17.643757

**Authors:** A. Chang, M. Peroš, A. Martashvili, S. Taherkhani, W. Magee, A. Claros, P.D. Dijkstra, S.G. Alvarado

## Abstract

An animal’s ability to adapt to a changing environment often requires the coordination of various traits. Across these traits, many covary with one another to generate a diversity of complex phenomes tuned to a given ecology. While many reports have documented trait covariation in populations, less is known about how plastic traits co-vary to facilitate adaptation in an individual. In African cichlids, morphology and behavior are two hallmarks driving the adaptive speciation of lineages within the East African Great Lakes. Here, we leverage social rank and body coloration as plastic model traits to understand the intersectional relationship shaping male competition in the African cichlid *Astatotilapia burtoni*. Addressing the need to disentangle the influence of environmental adaptation from social dynamics on color morphology, we conducted experiments rearing cichlids in visually distinct environments using blue and yellow gravel substrates to induce blue/yellow color morphs. Our results demonstrate that the visual environment significantly influences the emergence of male color morphs: yellow territorial males were more prevalent on brown gravel, whereas blue males predominantly appeared in blue backgrounds. Contrary to previous reports, we found that blue males consistently outcompete yellow males in direct contests. Furthermore, behavioral patterns changed over time, with blue males adjusting their aggression strategies based on their visual environment, while yellow males exhibited a higher propensity to flee. These findings indicate that animal coloration and behavior are intersectional plastic traits that interact to shape male competition and behavioral ecology. This study provides new insights into the dynamics of phenotypic plasticity, adaptive strategies in fluctuating environments, and trait covariation.

## INTRODUCTION

Phenotypic plasticity plays a crucial role in shaping how organisms react to a changing environment. This process can integrate changes in cellular/molecular substrates and environmental influences into adaptive traits, allowing for robust changes controlling one (or more) plastic trait(s). For example, triggering predatory cannibalism in drosophila larvae reared under crowded laboratory conditions can shape both the behavior and the morphology of mouthparts(Vijendravarma et al., 2013). Examples of phenotypic plasticity have been widely reported on single traits tied to metabolism, diet, and developmental timing across plants and animals (Iossa et al., 2019; Palmer et al., 2012; Sommer and Ogawa, 2011). However, little is known about the intersectional effects of plastic traits on one another. For example, an individual trait may present positive and/or negative feedback on other traits, resulting in complex phenomes better tuned to natural environmental changes than those typically observed in tightly controlled laboratory experiments.

Changes in behavior and animal coloration have various pleiotropic underlying physiologies(Engeszer et al., 2008; Lorin et al., 2018; Morris et al., 2001). For example, agouti signaling physiology in mice can drive feeding behavior and changes in pelage color(Carola et al., 2014; Moussa and Claycombe, 1999; Waterland and Jirtle, 2003). However, the study of multiple trait covariates has often been reported as descriptively modeling traits within a population, meriting a need for in-depth studies examining functional traits that directly influence an animal’s fitness. Here, we focus on two relevant, robustly plastic traits in an African cichlid to address this: (1) animal body coloration and (2) social behavior. The intersectional consideration of body coloration and behavior is fundamentally important across the animal kingdom and vital across various biological functions. For example, an animal’s body coloration can shape anxiolytic behaviors in the wild to remain cryptic(Culumber, 2016; Sih et al., 2004). While much is known about patterns of color variation at population scales(Hoekstra, 2006), less is known about repeated and reversible plastic responses of animal coloration that change throughout an individual’s life history(Alvarado, 2020).

Here, we focus on *A. burtoni* due to its robust and discrete behavioral and morphological differences concerning body coloration and social behavior. *A. burtoni* has blue and yellow body color phenotypes that are plastic to changes in visual and social environments(Fang et al., 2022; Fernald and Hirata, 1979; Korzan et al., 2008). Across both body color morphs, males can also become socially territorial (T) or non-territorial (NT)(Fernald and Hirata, 1977; Fernald and Hirata, 1979). T males are aggressive, claim territory, and court, whereas NT males do not claim territory and only attempt to court females in the absence of T males(Chen and Fernald, 2011; Maruska and Fernald, 2018). Within a community, T males are more aggressive and spend more time in agonistic interactions with males of the opposite color than NT males(Korzan and Fernald, 2006). While it has been shown that yellow T males typically outcompete blue T males in dyadic contests, the sampling conditions in identifying blue and yellow males have not been controlled for their social life history or rearing environments. However, current evidence suggests that blue-yellow polymorphism plays a role in male-male competition(Dijkstra et al., 2017; Korzan and Fernald, 2006; Korzan et al., 2008).

This prompts the following questions: (1) Does an individual’s social life history impact the ability of an individual to compete? And (2) Does an individual’s coloration shape its behavioral predispositions in social conflicts? By addressing these questions, we aim to clarify the relationship between social history, body coloration, and competitive behavior in *A. burtoni*. Understanding how these factors interact will provide deeper insights into the mechanisms of phenotypic plasticity and its role in shaping adaptive traits in dynamic environments. Furthermore, our findings will contribute to broader discussions on how plasticity influences social hierarchies, territoriality, and speciation in cichlids and other color-polymorphic species. Ultimately, disentangling the complex interactions between coloration and behavior will enhance our comprehension of how organisms navigate their ever-changing ecological landscapes.

## MATERIALS AND METHODS

### Animal husbandry and tissue collection

All experiments involving animals comply with relevant national and international guidelines, including but not limited to the Animal Welfare Act, the National Institutes of Health Guide for the Care and Use of Laboratory Animals, and the 3Rs Principle(Russell and Burch, 1960) (Replacement, Reduction, Refinement) under IACUC protocol #198 at Queens College. Fish were reared in communal settings in 118L tanks (76.2cm x 50.8cm x 30.5cm) under laboratory conditions that mimic their natural environment: pH 7.8 ± 8.2, 29°C, and 12-hour light/dark cycle with full-spectrum illumination(Fernald and Hirata, 1977). Water was chemically buffered using an automated sensor system (Neptune Systems Apex) to dispense salt (Seachem #0279) and water buffer (Seachem #0287). A brown gravel substrate closely approximated to the natural substrate (Pure Water Pebbles #30035) and halved terra-cotta pot (10cm dia x 10cm depth) shards were provided in all tanks. Pot shards allow territorial males to establish and maintain the territories necessary for successful reproductive behavior. All fish were fed daily at 9:00 A.M. ± 30 mins with cichlid pellets and flakes (Tetra AQ-77007)(Fernald, 1977). All males were collected within a few minutes of behavioral experiments. Fish were weighed using a digital scale (±0.001g) and measured for mouth-to-caudal trunk standard length before being sacrificed via rapid cervical transection.

### Experimental groups for behavioral analysis

#### Interactions between males of each color morph

To investigate all possible interactions among conspecifics, we picked non-territorial males who had never ascended before and displayed male-stereotype coloration(Fernald and Hirata, 1979). Tank gravel color has been shown to generate blue and yellow morphs of *A. burtoni(Dijkstra et al., 2024; Fang et al., 2022)*. Adult individuals were reared in tanks containing either blue gravel (Pure Water Pebbles #70021) or yellow gravel (Pure Water Pebbles #70111) with matching downwelling light from custom-made LED lighting set to a yellow or blue hue setting for four weeks. Induced blue (n=9) and yellow (n=10) experimental males were placed into the center compartment of a tank with one resident blue and one resident yellow on each side, separated by clear dividers. All males had three females and a terracotta pot in their compartments (Figure 1B). Tanks were filmed for 60 minutes starting at noon.

**Figure 1.**
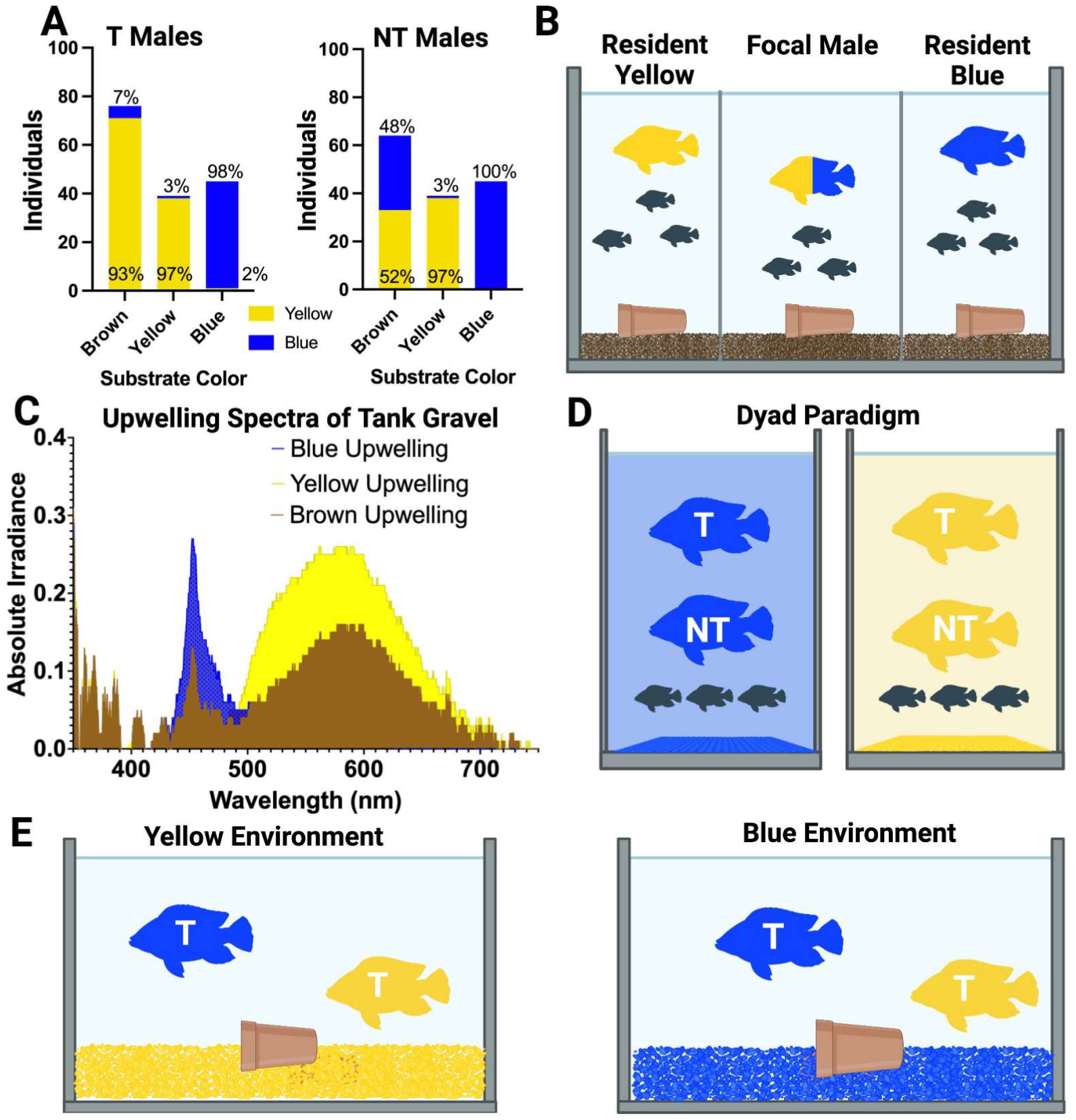
Experimental design for behavior analysis. **(A)** Ratio of blue to yellow morphs per substrate color **(B)** Social behavior assay investigating how T males interact **(C)** Spectrophotometer measures of the rearing environments in lab **(D)** Dyad paradigm consisting of two males and three females **(E)** Male competition assay

#### Dyad communities

To control for coloration and social history, adult individuals were placed into a dyad community containing two socially naive, size- and age-matched males and three females for four weeks to establish a social hierarchy, generating Territorial and Non-Territorial phenotypes (Figure 1D). To determine the effect of environmental color on social behavior, 40L tanks (50.8cm x 25.4cm x 30.5 cm) were supplied with yellow (n=17) or blue (n=18) gravel (Pure Water Pebbles #70021 and #70111). Each male pair was age and size-matched within 0.2mm. After the four-week induction, tanks were recorded starting at noon.

#### Male-male competition

To investigate if environment influences disputes between color morphs, we took territorial males of each color immediately after induction in a dyad and placed them in direct competition in either a yellow environment (n=6) or a blue environment (n=6) with a single territory (Figure IE). Males were size-matched as best as possible (mean difference 0.378 ± 0.204 cm) and placed in a tank with either blue or yellow gravel. Tanks were recorded for 3 hours to account for any changes in male tactics and to ensure the identification of a winner/loser. The winner was determined by which male occupied the territory at the end of the experiment.

### Behavioral Analysis and Scoring

All behaviors were recorded using video cameras (GoPro Hero8 Black) mounted on tripods (AmazonBasics, WT3540) for all video recordings. Tripods were positioned ~105cm away from the tank to capture all animals within the field of view. Video recordings were initiated at noon under “Standard” settings (1080p, 30FPS, linear) and manually scored after a 15-minute habituation period with BORIS event-logging software utilizing an ethogram for *A. burtoni* behavior (Table 1) (Friard and Gamba, 2016). All videos were scored blind to animal colors by setting hue to zero on LCD displays, making them black and white. Scorers could identify only sex-specific coloration, not body color (blue or yellow). Individuals who had unclear coloration or were out of view of the camera were excluded. Archival footage from Figure 1A was collected from videos of Dyad Communities done at Stanford University and Queens College CUNY on controlled blue/yellow/brown gravel substrates.

**Table 1.** Ethogram of *A. burtoni* behaviors.

| Behavior | Category | Description |
| --- | --- | --- |
| attack female | Reproductive | Snapping motion of the jaw of focal fish towards a female. |
| chase | Aggressive | Focal fish moves toward Fish B and follows as Fish B swims away without opening of jaw. |
| dig | Reproductive | Fish takes up the gravel in its mouth and expels it outside the territory. |
| pot entry/pot exit | Reproductive | Fish enters territory (the terracotta pot) |
| flee | Aversive | Focal fish quickly swims away from another fish in direct response to being followed. |
| forage | Other | Fish facing substrate or wall at an angle and pecking the area to eat. |
| lateral display | Aggressive | Focal fish adopts a lateral orientation, presenting its side to Fish B, often accompanied by fin extension and quivering movements. |
| lead | Reproductive | A male fish approaches a female, then reverses direction and guides the female toward the territory. The female follows, with the possibility of entering the territory. |
| quiver | Reproductive | A male fish performs rapid, tremulous movements towards another fish. |
| head to head | Aggressive | Two fish facing each other directly, often accompanied by operculum flaring. |
| bite | Aggressive | Snapping motion of the jaw of a focal fish towards Fish B |
| attack male | Aggressive | Snapping motion of the jaw of a male towards another male |

### Behavioral Transition Modeling

Scorelogs from videos were cleaned and organized in R and converted to CSV files. All models show the behaviors averaged amongst all trials. For the male-male competition experiments, we also computed matrices temporally per hour. We used Python version 3.12.3 and its libraries, Graphviz, pandas, and NumPy, to generate transition models to visualize the sequence of behaviors. The nodes represent each behavior with their size corresponding to the frequency the behavior occurred. Arrows represent the direction of transition and probability - higher probabilities are denoted by thicker arrows and lower probabilities are denoted by thinner arrows. Node positions were manually set to standardize each figure for clarity. All figures were completed using Biorender.

### Spectrophotometry

A portable spectrometer equipped with a fiber-optic collection cable (Ocean Insight) was used to measure the spectral profile in the tanks with brown, blue, and yellow gravel substrates (Figure 1C). Measurements were taken on the substrate (Supp Fig. 4A) and downwelling light from our LED system (Supp Fig 4B). Lighting was custom-made using RGB LED Flexible Light Strips (WenTop) housed in 2M long aluminum channels (Muzata).

### Statistical analysis

Score logs exported from BORIS software were analyzed in RStudio-2023.12.1-402, using base R and packages ‘ggplot2’ (version 3.4.4), ‘dplyr’ (version 1.1.3), ‘ggpubr’ (version 0.6.0), ‘tidyverse’ (version 2.0.0), ‘readr’ (version 2.1.4), ‘data.table’ (version 1.14.8) and ‘reshape2’ (version 1.4.4)(Gandrud, 2013; Kassambara, 2023; M, 2023; Team, 2024; Wickham, 2007; Wickham, 2016; Wickham et al., 2019; Wickham et al., 2023; Wickham et al., 2024). For the latency graphs, we constructed a data frame in base R that included behavior pairs, the time taken to transition from one behavior to another, the mean, and the standard error of the mean (SEM) for each behavior performed by each subject in each background color. Logs contained “Out of View” as a behavior, indicating periods where low visibility made it impossible to observe the fish’s actions. These instances were marked as state events with START/STOP logs and excluded from the transition time calculation. Independent t-tests with Benjamini-Hochberg corrections were then performed to compare fish color to background color, and point plots were generated to visualize the results. We used GraphPad Prism v. 10.3.1 to create figures and run the Fisher’s exact test, Wilcoxon-Mann-Whitney test, two-way ANOVA, and Repeated measures ANOVA.

## RESULTS

### The emergence of male color morphs is skewed by the visual environment

Male blue and yellow color morphs emerge naturally in the lab and the field(Fernald and Hirata, 1979). However, we noted that yellow males emerge more frequently in the visual environment. Using archival video footage of dyad paradigms taken on a brown substrate and full-spectrum white light, we noted that 93% of territorial males on brown gravel become yellow (p<0.0001), with non-territorial males split between blue (48%) or yellow (52%) morphs(Figure 1A). Similarly, we show that males reared in a blue or yellow visual environment (Figure 1D) almost entirely matched their visual background with the respective morph (Figure 1A, p<0.0001). Furthermore, when absolute irradiance was measured across brown/yellow/blue substrates, we noted that the spectral profiles of yellow and brown gravel were more similar than those of blue backgrounds (Figure 1C).

### Induced morphs interact with conspecifics differently within a community

In community environments (Figure 1B), induced focal yellow and blue males (center compartment) interact differently towards adjacent males belonging to each color morph. Blue males preferentially show aggressive displays towards conspecifics of the same color, while yellow displays indiscriminately (Figure 2A). Yellow focal males do more lateral displays (Mann Whitney U= 20, p= 0.0374) and attack opposing colors (Mann Whitney U=13, p= 0.0069). Blue males do more pot-related behaviors, shown by pot entries (U=15, p=0.0074) and exits (U= 16.5, p= 0.0137) than yellow counterparts (Figure 2A). To understand the possible motivations of induced blue and yellow males, we used behavior transition models of the focal male to assess the sequences of behavior qualitatively. From transition models, blue males were more likely to do reproductively connected behaviors in sequence (lead swim, pot entry/exit, and quivering at females) than yellow males (Figure 2BC). In contrast, no other differences were noted across aversive and aggressive behavior patterns (Figure 2BC).

**Figure 2.**
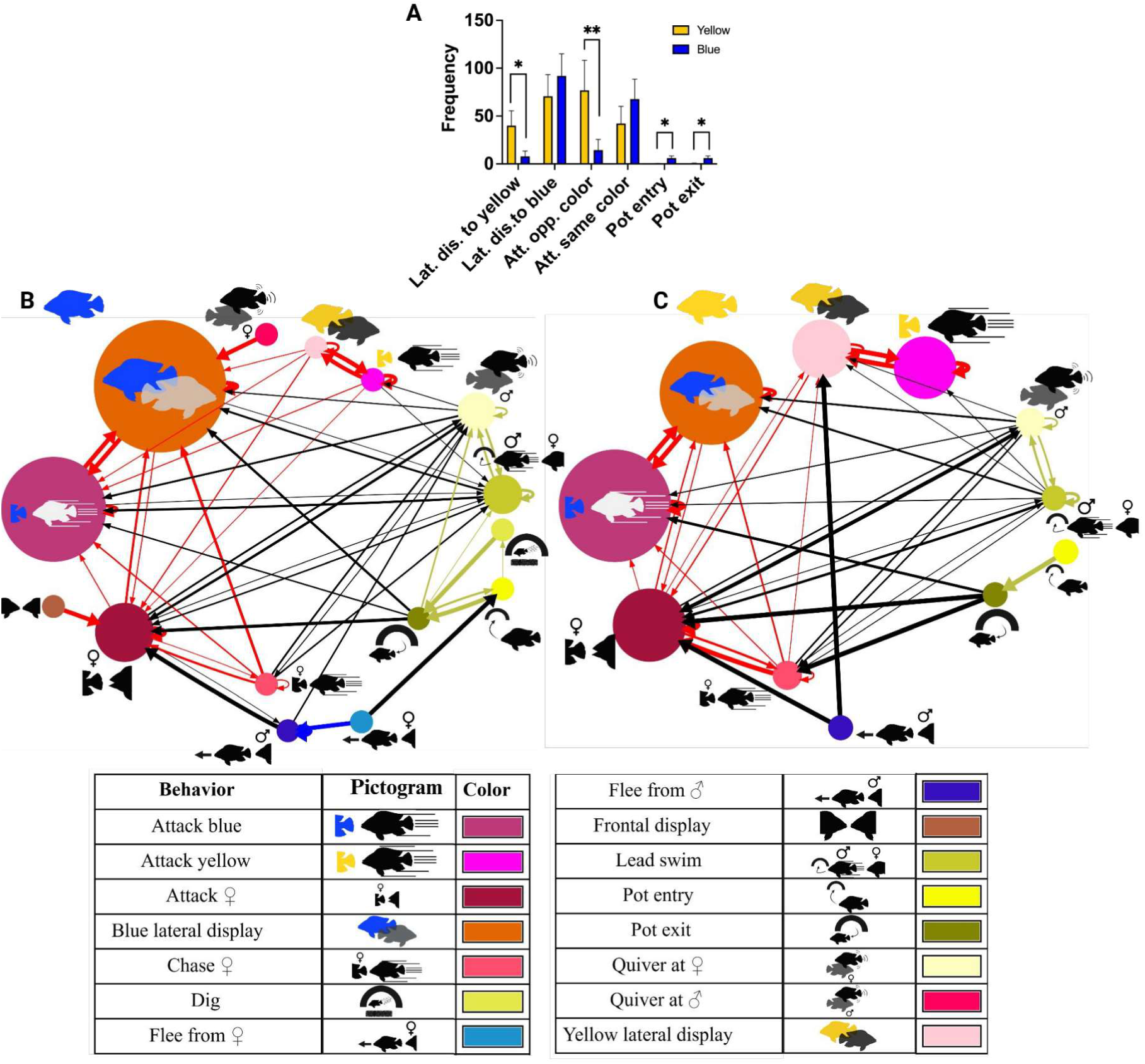
Blue and Yellow Males display different signatures of behavior **(A)** Behavior paradigm investigating how T males interact with each color morph. Total number of behaviors done by color-induced focal male (blue n=9, yellow n=10) when simultaneously exposed to both color morphs. Wilcoxon signed-rank test performed. * p<0.05, ** p<0.01. **(B-C)** Behavior transition models of T male behavior for each color morph. Probability of one behavior leading to another denoted by arrow thickness and frequency of behavior denoted by node size. The blue, red, and yellow arrows denote the category of behaviors, reproductive, aggressive, and aversive respectively. Black arrows represent links between behaviors of different categories. **(B)** Blue induced males **(C)** Yellow induced males

### Background coloration doesn’t affect social dynamics in a dyad paradigm

To investigate the intersectional role of color on social rank, we leveraged a dyad (Figure 1D) paradigm where we could control the colors of males and rank. This approach presented naturalistic manipulation of their coloration as lekking males(Fernald and Hirata, 1979) are more likely to reside in the same region on Lake Tanganyika with matching visual ecology(Horion et al., 2010). Both T and NT males showed stereotypical dyad behaviors with no marked differences due to the effect of background color (Figure S1). As described previously(Alcazar et al., 2014; Burmeister et al., 2005; Korzan et al., 2008; Maruska and Fernald, 2018; Maruska et al., 2011; Maruska et al., 2013), T males do significantly more chasing, biting behavior, and pot-residing than their NT counterparts (Figure 3). NT males flee significantly more (Figure 3E).

**Figure 3.**
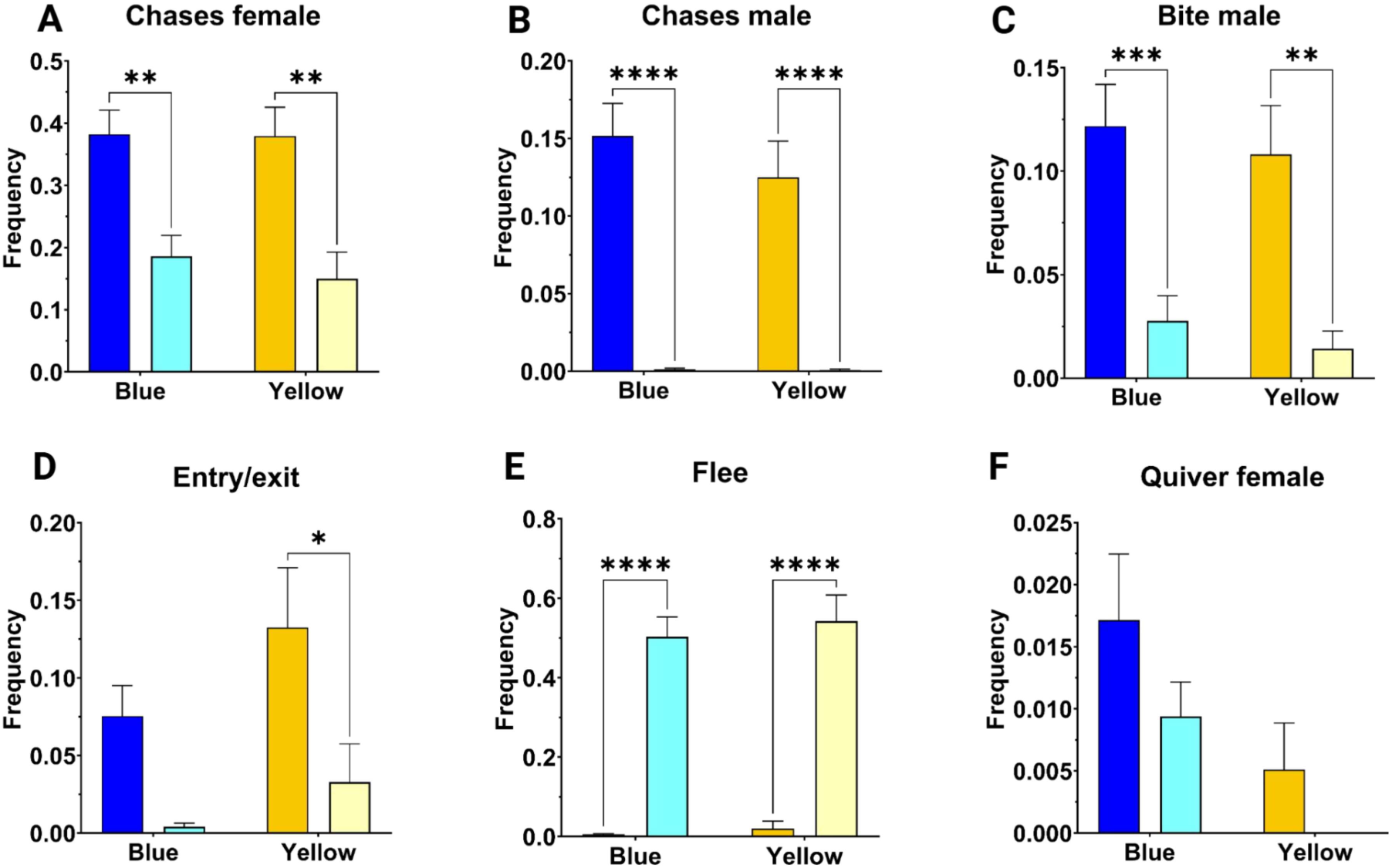
Social dyads show differences across rank but not color **(A-F)** Social behavior in blue and yellow male morphs in a social dyad paradigm. For all panels, blue (N=26) and yellow (N=18) male behavior is grouped by male social rank (T, darker bar; NT, lighter bar) with error bars representing standard error of the mean (SEM). Means between all comparisons were tested post-hoc of ANOVA using the Tukey multiple comparison test. Significance values reported as: * p<0.05; ** p<0.01; *** p<0.001; **** p<0.0001.

### Blue morphs outcompete yellow in a male-male competition assay

Previous work has reported yellow males are more aggressive and win in direct competition with blue fish(Korzan et al., 2008). To understand if environmental color shapes competition outcomes, we introduced two T males of each color from separate dyads in a tank with either yellow or blue substrate to contest over a terracotta pot (Figure 1E). In these contests, the blue fish outcompeted the yellow fish in 92% of trials regardless of the contest’s environmental condition. The two-way ANOVA, followed by Bonferroni’s multiple comparisons test, showed the effects of environmental color and fish coloration on behavior. The ANOVA revealed that in the yellow environment, fish behavior was responsible for 16.41% of the total variance (F_Behavior_(6,70)=4.903, p=0.0003; F_Color_(1,70)=6.281, p=0.0145, F_interaction_(6,70) = 12.26, p<0.0001). Blue fish showed significantly more chase (p= 0.0005) and pot entry/exit (p=0.0136) behaviors than yellow fish (Figure 4A). Yellow fish did significantly more flee behavior (p<0.0001) than blue. In the blue environment, behavior accounts for 13.07% of the total variance (Figure 4B, F_Behavior_(6,70) = 2.287, p= 0.0450; F_Interaction_(6,70) = 3.389, p= 0.0054). In the blue environment, yellow fish did significantly more flee behavior (p=0.0073).

**Figure 4.**
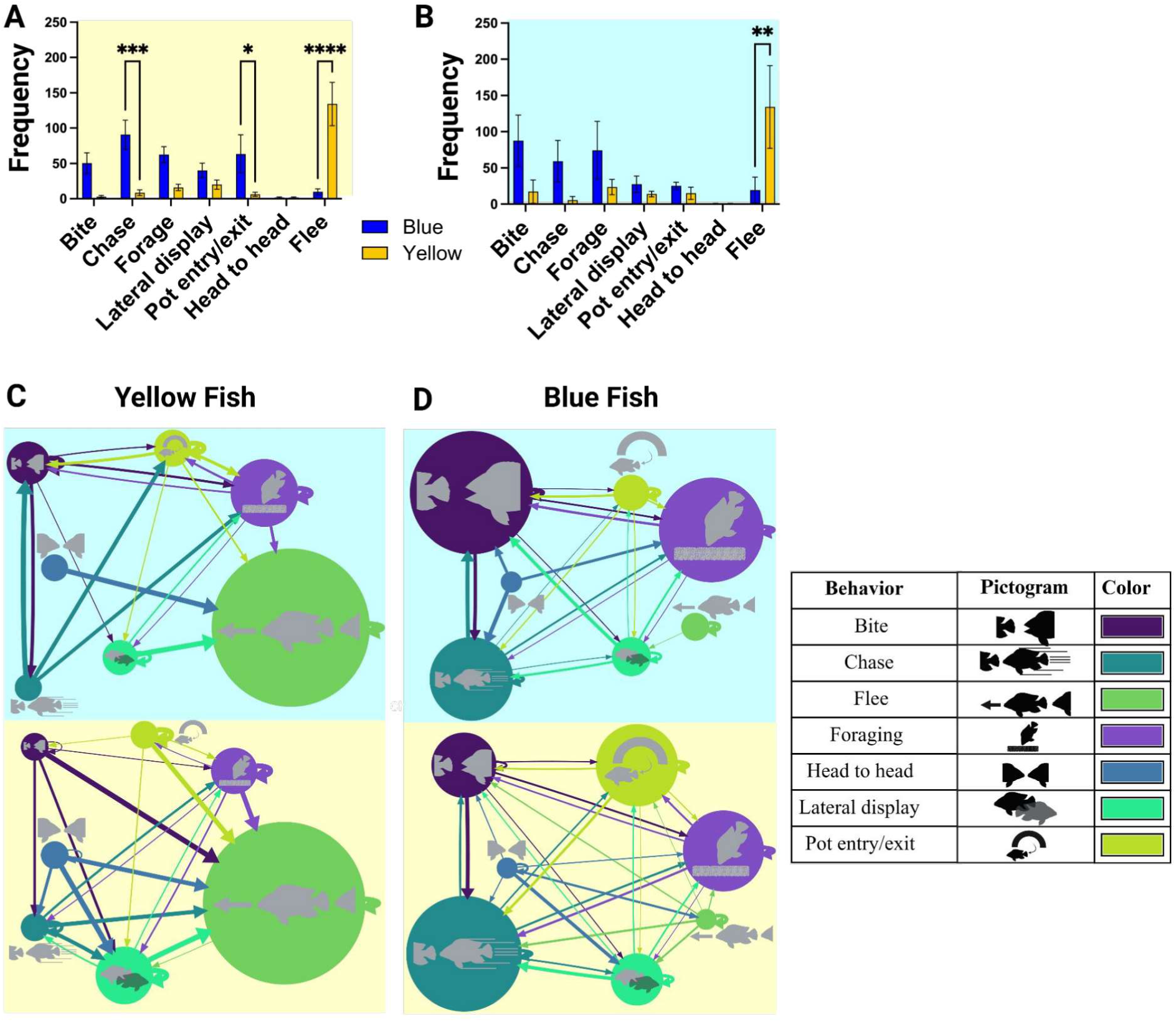
Blue and yellow males show different behavioral tactics dependent on environment **(A-B)** Behavior profiles of T males when in competing in **(A)** yellow environment and **(B)** blue environment. Significant comparisons using two-way ANOVA with Bonferroni multiple comparisons test are indicated: * p<0.05, ** p<0.01, *** p<0.001, ****p<0.0001. **(C-D)** Behavior transition models of yellow **(C)** and blue **(D)** fish in either environment during direct competition. Background color indicates test substrate color.

To investigate the temporal dynamics of the environment on the behavior of each color morph during the competition assay, we conducted repeated measures ANOVA followed by Tukey’s multiple comparisons test on the behaviors at each hour throughout the experiment. Blue fish implement different strategies for defending their territory depending on their surroundings. Of all the behaviors screened, bite, chase, pot entry/exit, and lateral display were significantly different and showed an interaction with environment and time. We see blue males interact between environment in the frequency of specific behaviors. The RM ANOVA revealed that the interaction between environment and time on bite, chase, pot entry/exit, and lateral display behaviors accounted for 11.9%, 10.1%, 4.52%, and 5.405% of the variance, respectively. In the yellow environment, blue fish do significantly more bite (p=0.0318), chase (p=0.0342), and pot entry/exit (p<0.05). The blue fish perform significantly more lateral display behavior in the first hour than in the second or third (Tukey post-hoc, p_1v2_=0.029; p_1v3_=0.0004) hours (Figure 5A). In a blue environment, blue fish perform significantly more lateral displays in the first hour compared to the second (p<0.05) and third (p<0.01) (Figure 5B).

**Figure 5.**
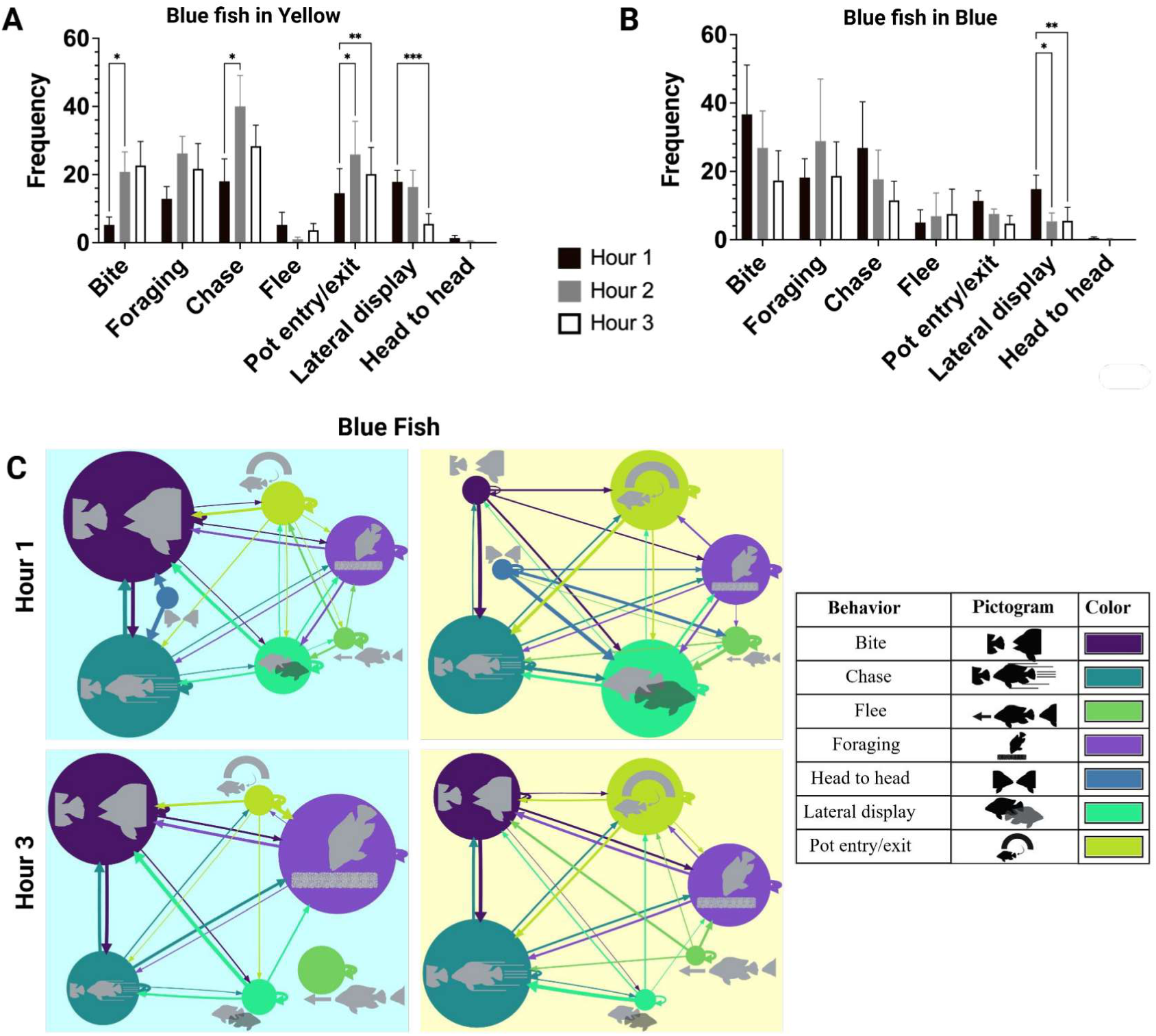
Temporal dynamics in behavior between blue/yellow males in a competition assay **(A-B)** Behavior profiles of blue T males competing in **(A)** yellow and **(B)** blue environments by hour of the most apparent behaviors related to territory acquisition and defense. Significant comparisons using two-way RM ANOVA with Bonferroni multiple comparisons test are indicated: * p<0.05, ** p<0.01, *** p<0.001. **(C)** Behavior transition models showing temporal differences in behavior between the start and end of the direct competition experiment. Background color indicates test substrate color.

We also measured the latency of the transitions between different behaviors to see if rearing color environment affected the decision-making of color morphs in a male competition assay. We noted a marked change in behavioral transition latencies of blue fish continuing “chase” to “chase” faster in a yellow background (M = 38.11, SEM = 2.84) compared to a blue background (M = 21.09, SEM = 2.99), t(247.84) = −4.13, p < .001, with an adjusted p-value of 0.0018. For the independent t-tests, some transitions had insufficient data to conduct the tests.

### Behavioral sequencing shows differences in blue/yellow male competition tactics

To understand the tentative motivations of differently colored male morphs during competition assays, we generated behavioral transition matrices to characterize sequences of behavior. We observed that yellow fish are more aggressive than blue fish in a yellow environment but tend to lose contests, as indicated by their tendency to flee (Figure 4E). In the third hour, yellow fish continued to do more aggressive behaviors, leading to fleeing (Supplemental figure 3C). In the yellow environment, blue fish initially lack clear behavioral patterns, but by the third hour, they become more aggressive, often chasing or biting after fleeing. In contrast, blue fish in a blue environment show a shift in behavior post-contest: they are initially offensive and revert to a neutral foraging behavior by the third hour. Additionally, in the yellow environment, blue fish display heightened aggression following a flee response at the beginning of the contest, a pattern not observed in the blue environment, with the frequency of aggressive behaviors declining by the third hour (Figure 5C).

## DISCUSSION

Our study shows that visual environment influences the emergence and rank-specific behavior of male color morphs in an African cichlid. First, we show that yellow territorial males appear more frequently on brown gravel. In contrast, blue males predominantly emerge in blue backgrounds, indicating that visual cues play a crucial role in morph determination. Second, color morphs exhibit distinct social behaviors, with blue males displaying aggression primarily towards their color and carrying out reproductively inclined behavior patterns with females. In contrast, yellow males show indiscriminate aggression. Third, in direct male-male competition, blue males consistently outcompete yellow males, contradicting previous findings that suggested yellow males were dominant(Korzan et al., 2008). Fourth, behavioral patterns shift over time, with blue males adapting their aggression strategies based on the environment, while yellow males continue aggressive displays but are more likely to flee. Finally, transition matrix analyses reveal that yellow males start aggressively but tend to retreat, whereas blue males strategically adjust their aggression, particularly in yellow environments, ultimately gaining dominance. Together, these data present evidence that animal coloration and behavior are intersectional plastic traits that can interact to shape male competition and behavioral ecology.

Changes in overall body coloration occur across various substrates and time scales. Rapid changes in pigmentation often rely on neural or physiological (or both) cues, as seen in cuttlefish(Hanlon and Messenger, 2018; Mthger et al., 2009). In contrast, gradual changes in pigmentation that comprise morphological color change result in changes in the density of pigment cells, their function, and gene transcription(Alvarado, 2020). In *A. burtoni,* neural and physiological substrates modulate body color and patterns. For example, its black eyebar changes rapidly through neurophysiological cues and during aggressive displays (Muske and Fernald, 1987a; Muske and Fernald, 1987b), whereas body coloration takes weeks to change via coordinated increases in pigment cell density and gene transcription(Fang et al., 2022). Our data reveal how plasticity in social behavior and body coloration can have reciprocal effects on one another. Previous work has shown that the melanocortin system contributes to aggression and coloration in cichlid fish(Dijkstra et al., 2017). Specifically, alpha-MSH administration is tied to the increased expression of yellow pigment in both morphs and aggression in only blue morphs. Conversely, administering agouti signaling peptide reduces aggression in both morphs but does not affect color. These effects have also been seen in reptiles, where different color morphs have been linked to aggression levels in agonistic interactions between conspecifics (Fernández et al., 2018). Previous studies on blue/yellow morphs have consistently shown that yellow morphs are more aggressive and outcompete blue morphs when placed in direct competition (Dijkstra et al., 2017; Dijkstra et al., 2024; Korzan and Fernald, 2006; Korzan et al., 2008). However, given the plastic nature of social status and body coloration, these studies did not control the environmental variables affecting these traits. These studies handpicked blue or yellow males from brown gravel substrates, which we show directly affects the likelihood of color morphs in subordinate male morphs (Figure 1A). Similarly, contests between males were completed by giving the individuals substantial habituation time within each other’s sight (between 4 to 30 days)(Korzan and Fernald, 2006; Korzan et al., 2008). Since *A. burtoni* rely heavily on visual cues for social information(Chen and Fernald, 2011; Grosenick et al., 2007), we suspect this may have skewed the outcomes of competitions between males of opposing colors. This could have been accentuated since socially naive yellow males in an open community setting show more threat displays than blue controls (Figure 2A). We also suspect that uncontrolled age matching in previous reports may have affected the fighting strategies employed between different males, which has been shown to shape the outcomes of male competition(Alcazar et al., 2014).

We also note that some of our data is partially supported by reports studying the aggressive and reproductive tendencies of blue/yellow males. While blue males typically lose in confrontations with yellow males, they show more aggression towards shoaling fish on brown gravel substrates(Korzan and Fernald, 2006). We show that blue male aggression leads to them winning in dyadic contests (Figure 4 and 5), but note that the environmental context can increase their aggression. For example, blue males on yellow backgrounds showed more chases, bites, and threat displays within hours 1-3 (Figures 4 and 5). We suspect the reported losses of blue males in previous studies may be attributable to their spectrally more yellow brown gravel substrates (Figure 1C), suggesting a “home team advantage” for yellow males. Our data also supports previous findings that blue males are more reproductively successful(Dijkstra et al., 2024) since they spend more time entering courting territories (Figure 2A) while indicating reproductive motivations through a higher likelihood of consecutive sexually reproductive behaviors (Figure 2BC). This is also supported by recent evidence showing that blue-rearing environments speed up the sexual maturity of blue versus yellow males(Moore et al., 2025).

We propose that systemic interactions between behavioral and color changes help males adapt to their shifting environment. In Lake Tanganyika, seasonal variations likely alter the visual ecology of the *A. burtoni* habitat(Horion et al., 2010). For instance, algal blooms at the lake’s southern tip can cause a dramatic shift toward green spectral dominance, making blue males more cryptic than their yellow counterparts while also increasing trophic activity in the water column. Conversely, the conspicuous yellow males are more easily detected by avian predators(Whitaker et al., 2021). Intriguingly, these color-based differences also correspond to variations in startle responses mediated by Mauthner neurons, with blue males exhibiting heightened startle readiness compared to yellow males on brown gravel. Additionally, shifts in yellow-blue body coloration are linked to social instability, where environmental changes increase the likelihood of males transitioning to yellow(Dijkstra et al., 2024).

Our findings underscore the intricate relationship between body coloration, social behavior, and competitive success in *A. burtoni*, demonstrating that these traits are not only plastic but also deeply interconnected. By revealing that visual environmental cues influence morph emergence, that color morphs exhibit distinct behavioral strategies, and that blue males outcompete yellow males in direct competition, our study challenges previous assumptions about dominance and aggression in this species. Furthermore, the behavioral plasticity exhibited by blue males in response to their environment suggests that social and ecological contexts play a significant role in shaping competitive outcomes. These insights highlight the importance of considering intersectional phenotypic plasticity when studying animal behavior and evolution. Future research should explore the underlying genetic and physiological mechanisms driving these plastic responses and assess their broader implications for social hierarchy formation, reproductive success, and species diversification in cichlids and other color-polymorphic species.

## ACKNOWLEDGEMENTS

We would like to thank funds provided by the Queens College Foundation, National Science Foundation (Award # 1921773), and The Professional Staff Congress of the City University of New York (TRADA-50-228) for funding provided for Annaliese Chang and Dr. Sebastian Alvarado. We would also like to thank the MARC-U-STAR program that funded Andrew Claros. Additionally, thank Dr. Maral Tajerian for revising this manuscript and providing constructive feedback on its writing.

