## Supplemental Figures for "Intersectional roles of trait covariation and phenotypic plasticity in the coloration and behavior of an African cichlid"

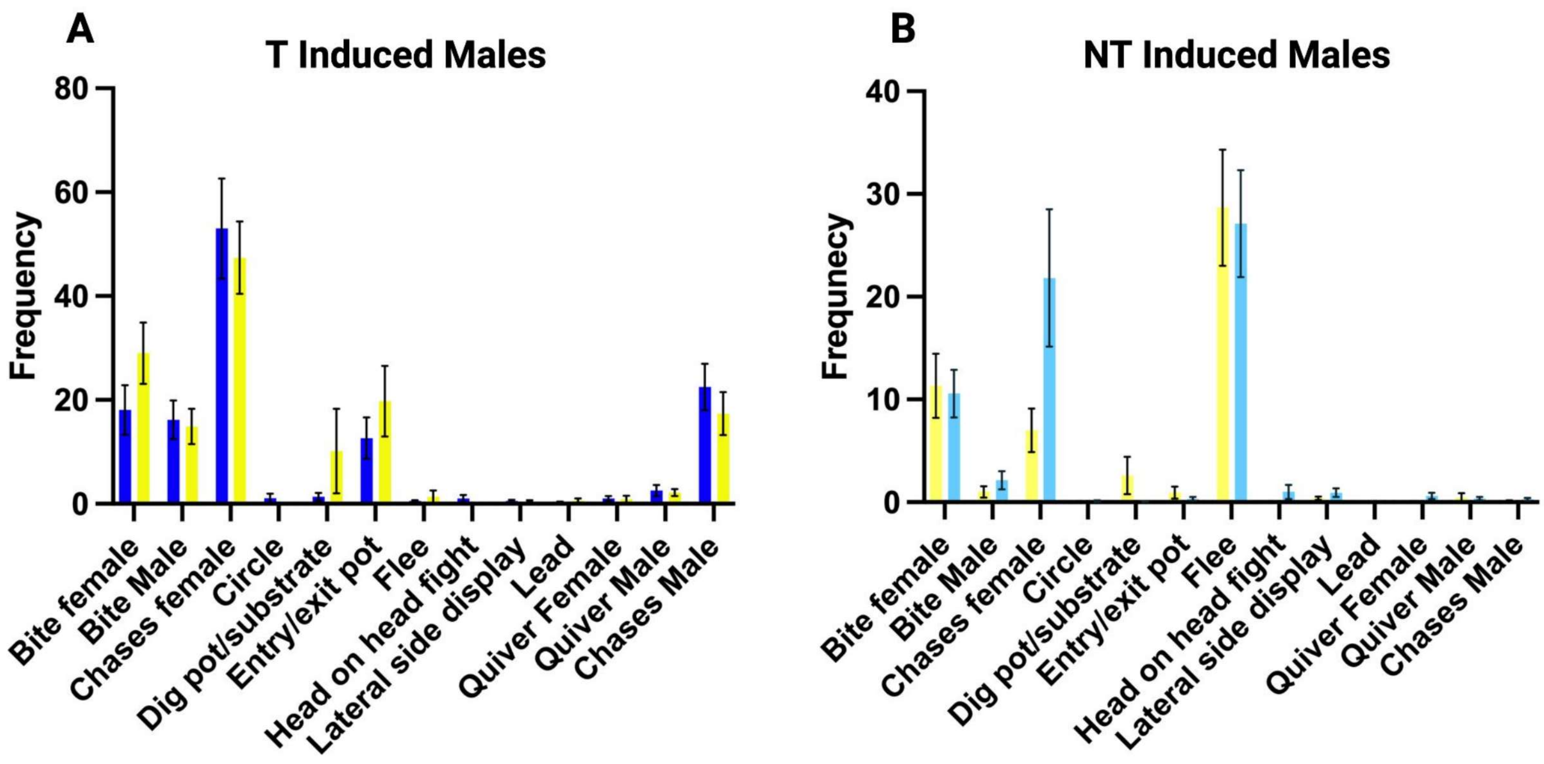


Supplemental Figure 1. Social rank behavior does not differ between color morphs (A) T males (B) NT male

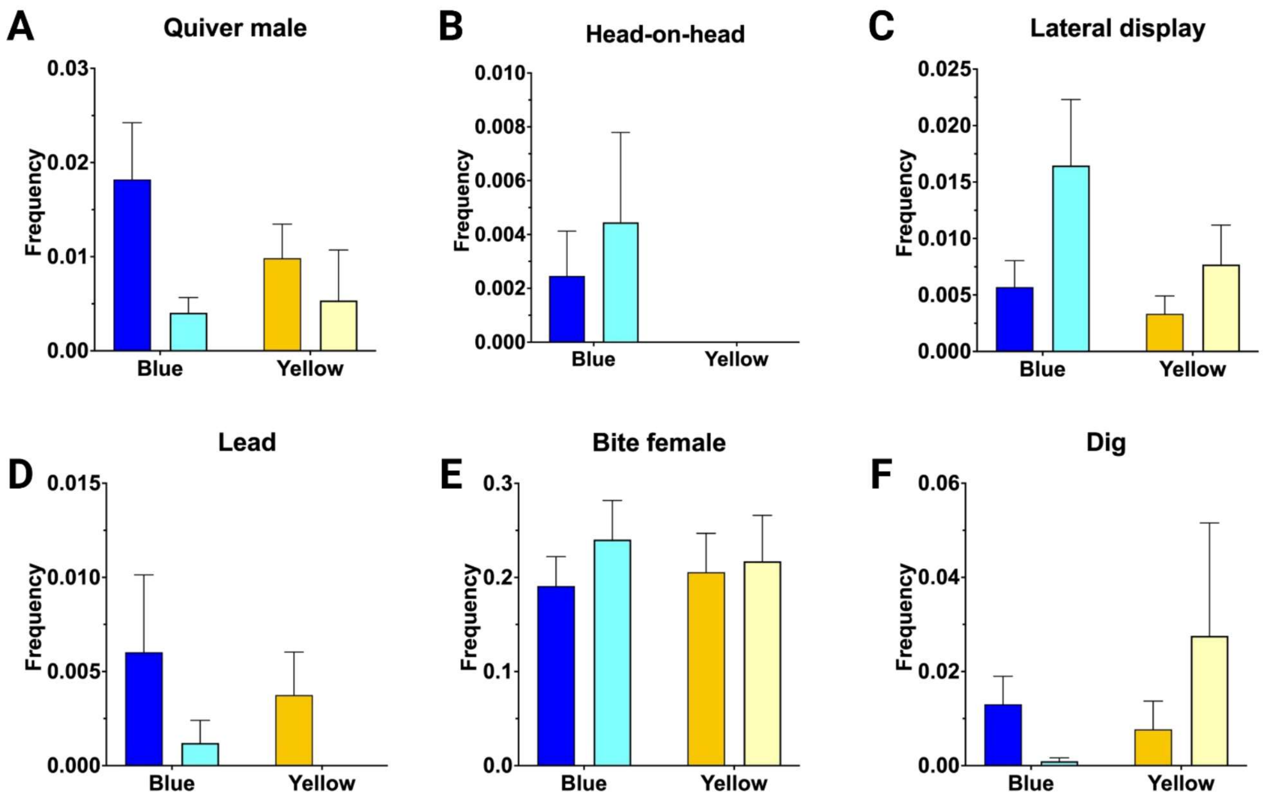


Supplemental Figure 2. Social behavior between blue and yellow male color morphs in a social dyad paradigm. For all panels, blue (N=26) and yellow (N=18) male behavior is grouped by male social rank (T, darker bar; NT, lighter bar) with error bars representing standard error of the mean (SEM). Means between all comparisons were tested post-hoc of ANOVA using the Tukey multiple comparison test.

| Transition | DF | t statistic | p value | Adjusted p value | MeanBlue±SEM | MeanYellow±SEM |
|---|---|---|---|---|---|---|
| Chase\|Chase | 247.839321 | -4.1286393 | 4.99E-05 | 0.001846198 | 21.0878±2.9924 | 38.1113±2.8367 |
| lateral display\|Bite | 33.0277877 | -2.8879877 | 0.00679232 | 0.125657868 | 19.4205±4.5662 | 50.9709±9.9247 |
| lateral display\|Foraging | 51.452446 | -2.3557975 | 0.02233026 | 0.275406496 | 27.1612±8.7819 | 70.1725±16.0068 |
| pot entry/exit\|Bite | 33.3834246 | 2.20571193 | 0.03439451 | 0.318149216 | 31.4091±8.2581 | 12.5275±2.2543 |
| Foraging\|lateral display | 33.4830233 | -1.7793078 | 0.08426894 | 0.519658451 | 32.4703±6.331 | 60.1514±14.2108 |
| Bite\|Foraging | 127.237799 | 1.78826546 | 0.07611303 | 0.519658451 | 48.3284±5.7766 | 34.0964±5.4744 |
| Chase\|lateral display | 41.9572099 | -0.6148348 | 0.54198414 | 0.752499684 | 27.1878±12.8781 | 36.5954±8.2628 |
| Foraging\|Bite | 135.786751 | 1.37684554 | 0.17082581 | 0.752499684 | 47.7113±10.9417 | 30.9866±5.2757 |
| Bite\|Chase | 281.919106 | -0.6980872 | 0.48569807 | 0.752499684 | 11.9696±2.2192 | 14.1768±2.2522 |
| Bite\|Bite | 210.37469 | 0.77207258 | 0.44093789 | 0.752499684 | 37.4182±4.482 | 32.8702±3.8224 |
| Foraging\|Chase | 52.914376 | 0.84428492 | 0.40231124 | 0.752499684 | 22.6842±4.8429 | 18.1193±2.4041 |
| Bite\|lateral display | 41.8439221 | -0.8216954 | 0.41590774 | 0.752499684 | 10.1436±2.7246 | 14.6172±4.7135 |
| pot entry/exit\|pot entry/exit | 70.7182369 | 0.79555357 | 0.42895383 | 0.752499684 | 29.2892±13.792 | 18.0369±3.1357 |
| pot entry/exit\|lateral display | 14.207311 | 1.33937514 | 0.20148249 | 0.752499684 | 38.9145±14.2065 | 19.0417±4.2803 |
| Chase\|head to head | 1.07014255 | -0.8824855 | 0.53176063 | 0.752499684 | 19.6285±6.6285 | 51.388±35.373 |
| pot entry/exit\|Chase | 28.5910968 | -1.1289726 | 0.26829335 | 0.752499684 | 7.4938±2.3711 | 10.6065±1.407 |
| Chase\|pot entry/exit | 20.4880682 | 0.67247528 | 0.50879508 | 0.752499684 | 33.9964±10.7258 | 26.4349±3.375 |
| pot entry/exit\|Foraging | 42.6128544 | 0.65665788 | 0.51493171 | 0.752499684 | 56.4596±20.0125 | 39.8464±15.4782 |
| Foraging\|pot entry/exit | 34.2045229 | 0.60507714 | 0.54912139 | 0.752499684 | 39.8928±12.0789 | 31.2116±7.7422 |
| lateral display\|Flee | 12.867413 | 0.67559431 | 0.51125772 | 0.752499684 | 72.6861±30.6061 | 45.9794±25.0189 |
| Flee\|Bite | 1.09519429 | 0.90211913 | 0.52193605 | 0.752499684 | 32.987±12.227 | 21.7021±2.6424 |
| Flee\|lateral display | 7.43507605 | 1.23622549 | 0.25400026 | 0.752499684 | 183.7547±66.3633 | 92.4518±32.4136 |
| Flee\|Flee | 32.1197797 | 1.25873067 | 0.21719982 | 0.752499684 | 79.3932±7.86 | 59.1195±14.0585 |
| Flee\|Foraging | 3.69152208 | 0.91250112 | 0.417126 | 0.752499684 | 150.0773±60.0027 | 85.2525±38.0325 |
| Flee\|pot entry/exit | 5.7153788 | 0.66523398 | 0.53180994 | 0.752499684 | 94.3168±40.4244 | 58.4283±35.7257 |
| pot entry/exit\|Flee | 2.02121157 | -1.9590128 | 0.1878594 | 0.752499684 | 24.128±12.3674 | 357.856±169.9057 |
| Bite\|Flee | 1.00827524 | 1.29176108 | 0.41807119 | 0.752499684 | 156.6095±97.3815 | 30.5562±6.2597 |
| Chase\|Flee | 2.12907716 | 0.62579036 | 0.59192979 | 0.782192935 | 38.102±20.2116 | 25.2538±3.6082 |
| lateral display\|lateral display | 95.5049159 | -0.5028469 | 0.61622958 | 0.786223952 | 48.3531±11.0341 | 56.2111±11.0661 |
| Chase\|Bite | 238.840205 | -0.4506792 | 0.65262962 | 0.804909861 | 28.8961±3.1456 | 30.6853±2.4219 |
| Bite\|pot entry/exit | 61.4734323 | -0.3955306 | 0.69382063 | 0.828108493 | 28.9625±7.9194 | 33.2006±7.2174 |
| Chase\|Foraging | 139.674309 | -0.3607674 | 0.71881837 | 0.831133739 | 41.724±6.7853 | 45.1012±6.449 |
| lateral display\|Chase | 64.1683744 | -0.3035546 | 0.76244934 | 0.853960768 | 32.4685±6.3909 | 34.9412±5.051 |
| Foraging\|Flee | 1.16527542 | 0.34030498 | 0.78472071 | 0.853960768 | 84.398±67.398 | 60.5633±19.0523 |
| lateral display\|pot entry/exit | 18.834197 | -0.1979629 | 0.84519508 | 0.893491943 | 41.422±16.1036 | 45.2149±10.3814 |
| Foraging\|Foraging | 372.941918 | -0.0889596 | 0.92916173 | 0.954971782 | 32.7974±4.7781 | 33.3742±4.3814 |
| Flee\|Chase | 1.17133361 | -0.0613562 | 0.95983884 | 0.959838838 | 36.437±25.579 | 38.07±7.3537 |
| head to head\|Chase | NA | NA | NA | NA | 9.3765±4.1235 | 33.5 |
| lateral display\|head to head | NA | NA | NA | NA | 125.203 | 159.077±128.9924 |
| head to head\|Bite | NA | NA | NA | NA | 2.756 | 86.063 |
| Foraging\|head to head | NA | NA | NA | NA | 29.528 | 76.092 |
| head to head\|Foraging | NA | NA | NA | NA | 51.979 | 15.029 |
| head to head\|lateral display | NA | NA | NA | NA | NA | 74.9203±68.7651 |
| head to head\|Flee | NA | NA | NA | NA | NA | 20.2285±4.2285 |
| Flee\|head to head | NA | NA | NA | NA | NA | 226.464±98.026 |
| head to head\|head to head | NA | NA | NA | NA | NA | 79.5035±28.2445 |

Supplemental Table 1. Independent t-tests with Benjamini-Hochberg corrections completed to compare latency between behaviors for blue fish in each environment. Red indicates significant result.

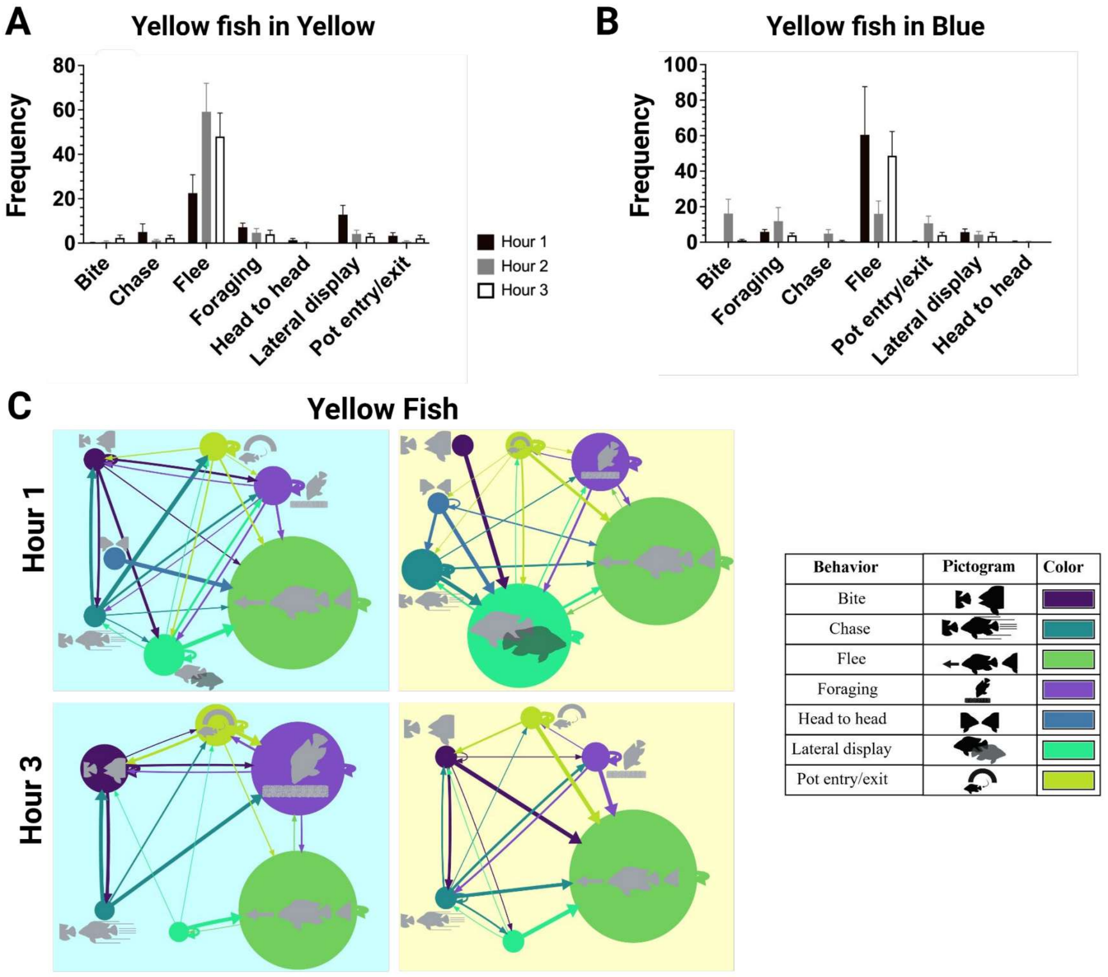


Supplemental Figure 3. Temporal dynamics of yellow fish during competition assay in **(A)** yellow and **(B)** blue environment by hour of the most apparent behaviors related to territory acquisition and defense. **(C)** Behavior transition models showing temporal differences in behavior between the start and end of the direct competition experiment. Background color indicates test substrate color.

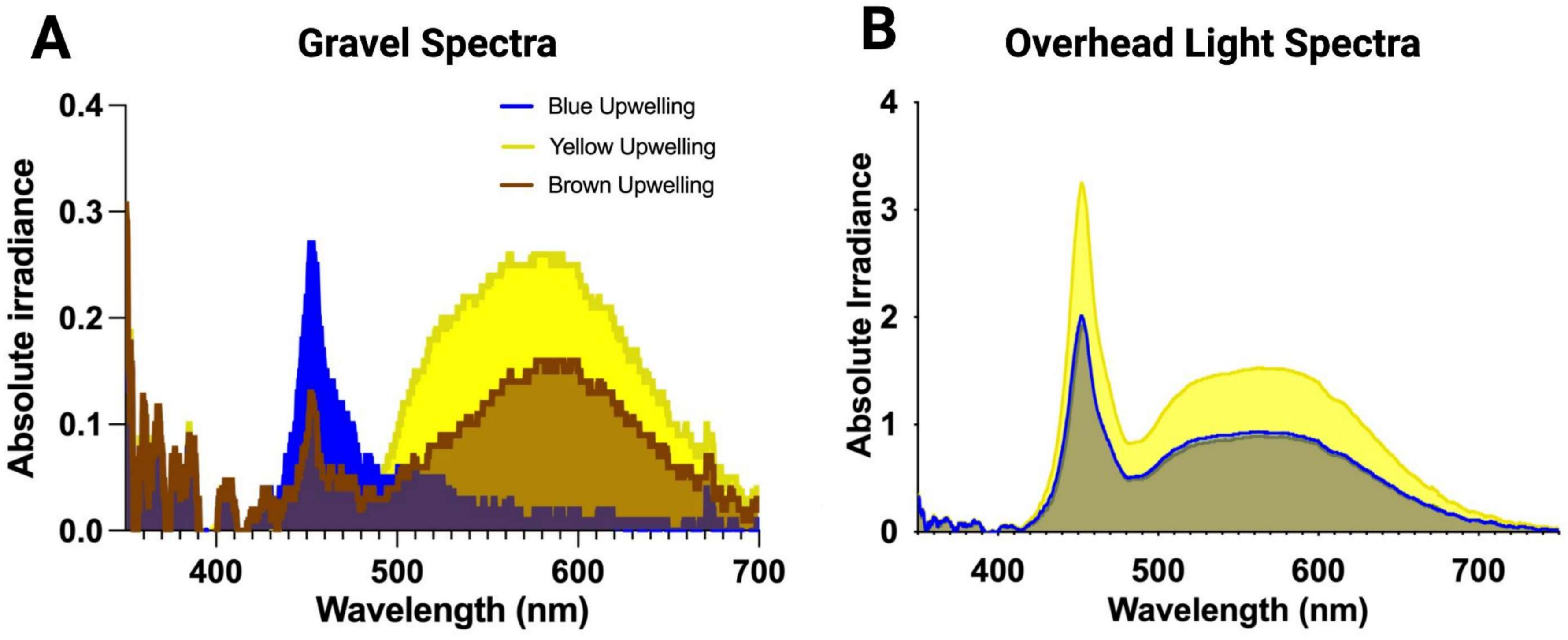


Supplemental Figure 4. Spectrophotometer readings of lighting
